# Specific proteolysis mediated by a p97-directed proteolysis-targeting chimera (PROTAC)

**DOI:** 10.1101/2024.03.08.584142

**Authors:** Constanza Salinas-Rebolledo, Javier Blesa, Guillermo Valenzuela-Nieto, David Schwefel, Natalia López González del Rey, Maxs Méndez-Ruette, Janine Burkhalter, Luis Federico Bátiz, Ronald Jara, José A. Obeso, Pedro Chana-Cuevas, Gopal P. Sapkota, Alejandro Rojas-Fernandez

## Abstract

The p97 protein is a member of the AAA + family of ATPases. It is a mechanoenzyme that uses energy from ATP hydrolysis to promote protein unfolding and segregation actively. The unfolded products of p97 are presented to the 26S Proteasome for degradation. p97 substrate recognition is mediated by adaptors, which interact with substrates directly or indirectly through ubiquitin modifications, resulting in substrate funnelling into the central pore of the p97 hexamer and unfolding. We have engineered synthetic adaptors to target specific substrates to p97, using the extraordinary intracellular binding capabilities of camelid nanobodies fused to the UBX domain of the p97 Adapter FAF1. In such a way, we created a p97-directed proteolysis-targeting chimera (PROTAC), representing a unique E3 ubiquitin ligase-independent strategy to promote specific proteolysis.

## Introduction

The ubiquitin-proteasome system (UPS) regulates protein abundance by specific E3 ubiquitin ligases, which catalyze ubiquitin chain formation on the substrates, inducing their proteasome-mediated degradation^1-4^. The UPS as an efficient natural negative-regulatory mechanism has inspired the development of the Proteolysis-Targeting Chimera (PROTAC) technology, involving synthetic heterobifunctional molecules able to recruit a protein of interest (POI) to a ubiquitin E3 ligase to induce its proteasomal degradation^5-16,17,18^

The nature of PROTACs varies from small molecules to protein domains and antibody fragments. Several camelid Nanobodies are capable of binding intracellular target proteins selectively, with a high affinity. When Nanobodies are overexpressed in cells, they are also known as intrabodies^15,19,20^. Nanobodies fused to ubiquitin E3 ligase substrate receptors trigger protein degradation of ectopic and endogenous targets. For instance, the AdPROM system consisting of a fusion of the E3 substrate receptor VHL with nanobodies against a range of POIs leads to the efficient degradation of endogenous POI targets^8,9,20-24^. Nanobodies have also been engineered and fused directly to the active domains of Ubiquitin E3 ligases, such as the antibody RING-mediated destruction system (ARMeD), which uses the RING finger domain of the Ubiquitin E3 ligase RNF4 fused to a nanobody. A valuable feature of the ARMeD system is its independence of the endogenous ubiquitin E3 ligases^15^.

In addition to conventional ubiquitination and proteasomal-mediated degradation, the AAA-type ATPase p97 assists the proteasome in the specific selection of substrate degradation in eukaryotic cells^25^. The key mechanism of action of p97 is the disassembly of protein complexes^26^ through its ATP-dependent *segregase* and *unfoldase* activity^27-29^. p97 uses ATP hydrolysis as a source of energy to ‘segregate’ ubiquitylated protein complex subunits from their binding partners, or even for the unfolding of protein aggregates^30,31^, such as tau amyloid fibers^32^ and Huntingtin Exon 1^33^. This action is mediated by two types of adapters: the UBX-like domain (UBX-L, also known as the “ubiquitin-binding domain” [UBD]) adapters such as Ufd1, NLP4, p47, FAF1, SAKS, UBXD7 and UBXD8, and the UBX-only adapters, such as p37, UBXD1, UBXD2, UBXD3, UBXD4, UBXD5, UBXD6, VCIP135 and YOD1^34-38^. p97 regulates ER-mitochondrial association by disassembling Mfn2 complexes upon PINK/Parkin phosphoubiquitination^39^ and the mitochondrial extraction of MARCH5^40^. It also co-localizes with protein aggregates involved in several neurodegenerative diseases, suggesting that p97 *segregase* and *unfoldase* activities could be related to the proteolytic control of proteins of therapeutic interest^41-46^. For instance, p97 is recruited to poly-Q aggregates *in vitro^31^* and to inclusion-positive neurons in Huntington’s disease patients^42^. Abnormal protein aggregation is observed in several pathologies such as body myopathy associated with Paget’s disease, frontotemporal dementia^47,48^, Charcot–Marie–Tooth disease^49^, amyotrophic lateral sclerosis^50^, and Parkinson’s disease^46^. Ubiquitin is often found as a resident protein within aggregates related to neurodegenerative diseases suggesting potential dysfunction of ubiquitin-mediated degradation signaling^51-54^.

Motivated by these characteristics, we engineered a synthetic p97 adapter by fusing the UBX domain of the FAF1 protein to camelid nanobodies, to assemble a novel p97-based PROTAC technology (**p97-PROTAC**). This new chimera efficiently targets proteins for segregation and proteasome-mediated degradation in a ubiquitin-independent manner.

## Results

### p97-PROTAC subcellular recruitment and activity in cells

p97 adapters and cofactors lead to the specific recognition of ubiquitin-modified and unmodified substrates along with simultaneous entry into the central pore of p97 hexamers, followed by unfolding and subsequent proteasomal degradation^55,56^. We hypothesized that p97 adapters could be engineered to target non-natural substrates of clinical interest for degradation. Thus, we generated a synthetic chimera consisting of the Ubx domain of the p97 adapter FAF1 fused via a linker to a GFP-specific camelid nanobody (Ubx-Nb^(GFP)^), which is capable of recognizing both GFP and YFP tagged proteins^57^, to yield a novel PROTAC based on p97 activity (Figure 1A). Using fluorescence microscopy, we then evaluated if the Ubx-Nb^(GFP)^ recognizes GFP-fusion proteins with different sub-nuclear localization by analyzing Ubx-Nb^(GFP)^ recruitment to a group of differentially located targets: i) GFP-Coilin (Nuclear and Cajal Bodies), ii) GFP-Emerin (a type II integral membrane protein residing principally at the inner nuclear membrane), and ii) GFP-ETV1 (Nuclear transcription factor) in HeLa cells. We observed efficient Ubx-Nb^(GFP)^ recruitment to diverse nuclear locations (Figure 1B).

**Figure 1.**
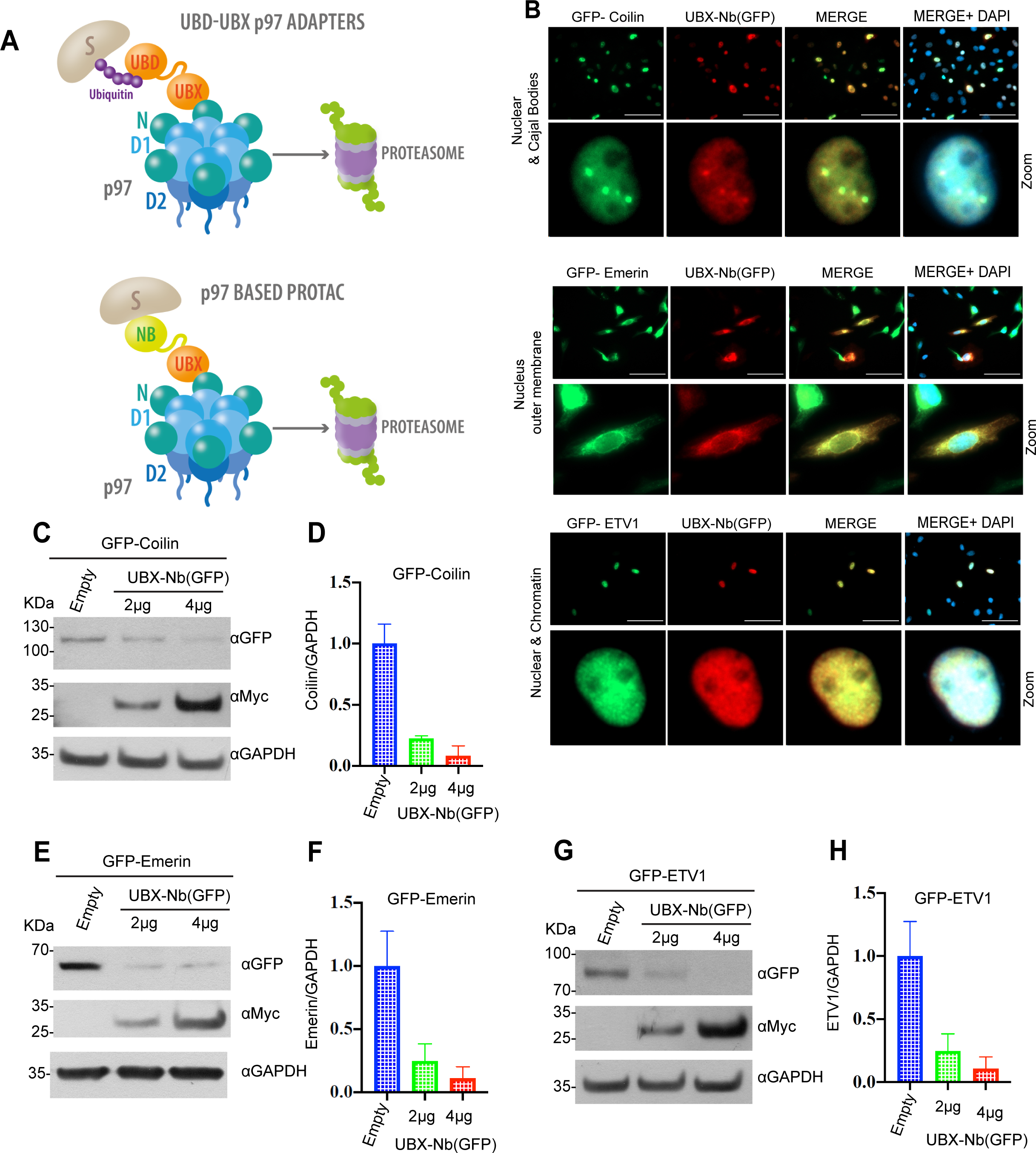
p97-mediated proteasome degradation. **A)** Diagram of p97 presenting ubiquitinated proteins to the proteasome via a UBX domain contacting adaptor (Top). p97-PROTAC system, consisting of a UBX domain fused to a nanobody (Nb) that recruits substrates for p97-mediated segregation, unfolding and proteasomal mediated degradation (Bottom). **B)** p97-PROTAC (UBX-Nb^(GFP)^) recognizes GFP tagged proteins at different cellular locations. HeLa cells were seeded on coverslip and co-transfected with UBX-Nb^(GFP)^ and GFP-Colin, GFP-Emerin and GFP-ETV1. Cells were fixed and we performed immunofluorescence for anti-myc tag to verify the expression of UBX-Nb^(GFP)^ and evaluated the colocalization. **C)** Western blot analysis of GFP-Coilin degradation by transfection with p97-PROTAC (UBX-Nb^(GFP)^), **D)** quantification of C;. **E)** Western blot analysis GFP-Emerin degradation by transfection with p97-PROTAC (UBX-Nb^(GFP)^), **F)** quantification of E;. **G)** Western blot analysis GFP-ETV1 degradation by transfection with p97-PROTAC (UBX-Nb^(GFP)^), **D)** quantification of G; Western blots were quantified and statistically analyzed using a student’s t-test. P < 0.05 compared to controls. n=3.

Next, we studied the effect of Myc-tagged Ubx-Nb^(GFP)^ PROTAC expression on the levels of GFP-Coilin, GFP-Emerin and GFP-ETV1. We observed a significant reduction in the levels of all GFP target proteins with increasing amounts of Myc-Ubx-Nb^(GFP)^, indicating that the p97-PROTAC - Ubx-Nb^(GFP)^, is sufficient to trigger specific proteolysis of target proteins (Figure 1C-1H). Consistently, no degradation was observed when the GFP nanobody was replaced by a negative control nanobody (Supplementary Figure 1A-C). Therefore, this suggested that p97-PROTAC could be an alternative targeted protein degradation (TPD) approach to the conventional ubiquitin ligase-based PROTACs and might be useful for the degradation of large protein complexes,integral membrane protein at the inner nuclear membrane and toxic aggregates due to its *unfoldase* and *segregase* activities.

### Specific degradation of proteins within liquid-liquid phase separation structures

We sought to test the efficacy of the p97-PROTAC system for the degradation of proteins whose expression was driven by endogenous promoters. To further interrogate the p97-PROTAC for its ability to induce degradation of proteins within aggregates and packed in highly condensed regions, we chose *53BP1* as the target protein, as it is a well-known component of liquid-liquid phase separation structures (LLPS)^58,59^.

First, we generated a knock-in cell line by fusing a yellow fluorescent protein to the N-terminus of the endogenous 53BP1 protein using CRISPR/Cas9 technology. 24h after transfection with a forward and reverse gRNA and the donor vector, Cas9 D10A nickase was induced with doxycycline. Single YFP-positive cell clones were isolated by cell sorting (Figure 2A). Natural DNA damage that occurs during DNA replication induces the accumulation of 53BP1 Fluorescence signal in distinct nuclear foci. The foci are typical for 53BP1, which has been described to accumulate in nuclear bodies during the G1 phase of the cell cycle^60^. The YFP-53BP1 Knock-In (KI) clones were analyzed by fluorescence microscopy and a heterozygote clone was selected (Figure 2B). We characterized 53BP1 foci in U2OS YFP-53BP1 (KI) cells by structured illumination microscopy (SIM) (Figure 2C). Further, we evaluated the localization of the YFP-53BP1 KI and endogenous 53BP1 by SIM microscopy and demonstrated the expected location for the knock-in fusion (Figure 2D). Accordingly, the YFP-53BP1(KI) cell line fully recapitulated the localization of endogenous 53BP1 and the endogenous 53BP1 promoter controls the YFP-53BP1 KI expression. 53BP1 nuclear bodies have been described as membrane-less organelles, organized by liquid-liquid phase separation (LLPS)^58,59^. Hence, we used the YFP-53BP1 KI cells as a model to study the recruitment and function of the p97-PROTAC Ubx-Nb^(GFP)^ to target LLPS. We observed that the Ubx-Nb^(GFP)^ construct was successfully recruited to nuclear foci characterized by the 53BP1 location (Figure 2E). Importantly, Ubx-Nb^(GFP)^ expression significantly reduced 53BP1 levels, suggesting that our p97-PROTAC technology can specifically trigger the degradation of proteins within LLPS compartments (Figure 2F & 2G).

**Figure 2.**
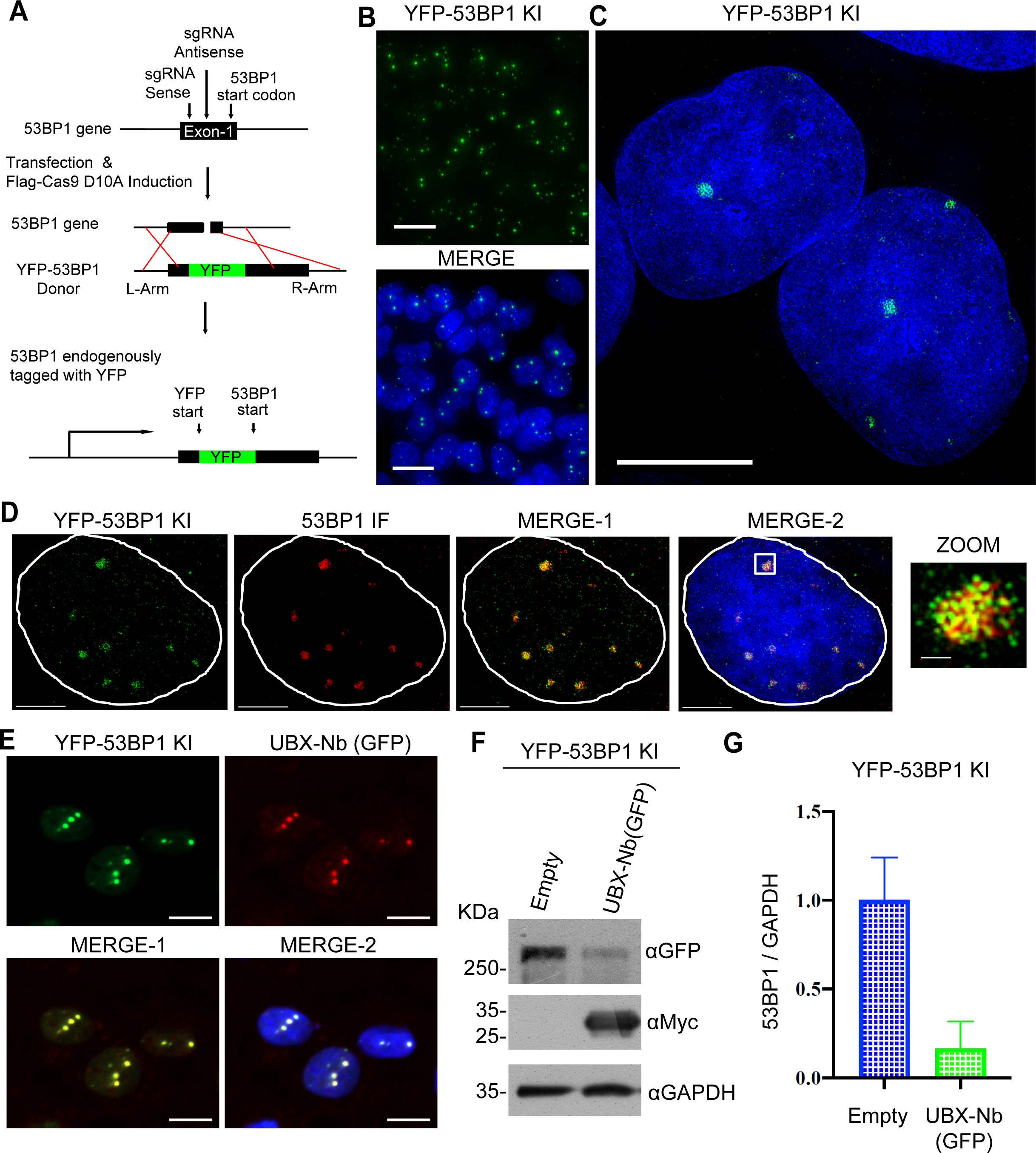
Targeting Liquid-liquid phase separation proteins by a p97-PROTAC. **(A)** Strategy for inserting a YFP tag on the N-terminus of the 53BP1 gene in U2OS SEC-C cells using CRISPR/Cas9 D10A. **B)** Selected Knock-In (KI) YFP-53BP1 clones isolated via flow cytometry. Clones were confirmed via florescence microscopy (GE Deltavision Widefield). **C)** Super-resolution images obtained with a Delta Vision OMX V4 structured illumination microscope (3D-SIM). **D)** Immunofluorescence against 53BP1 (red) and colocalization with YFP-53BP1 in Knock-In cells. Images were obtained using a Delta Vision OMX V4 structured illumination microscope (3D-SIM). **E)** Recruitment of the p97-PROTAC UBX-Nb^(GFP)^ (Red) to YFP-53BP1 (green) within Liquid-liquid phase separation structures. Data were obtained with a High-content Celldiscoverer 7. UBX-Nb^(GFP)^ was detected using its myc-tag. **F)** Western blot analysis YFP-53BP1 degradation by UBX-Nb^(GFP)^ transfection in the knock-In U2OS cells, **G)** Quantification of F. Western blots were quantified and statistically analyzed using a student’s t-test. P < 0.05 compared to controls. n=3.

### Endogenous p97 expression in the brain

Conventional PROTAC technologies exploit the activity of E3 ubiquitin ligases and accordingly require the expression of a suitable ligase to cause target ubiquitination and degradation. Likewise, the p97-PROTAC function requires the endogenous p97 expression. Therefore, we studied the expression of p97 in the brains of animal models. We observed ubiquitous expression in the cortex of non-human primates (NHPs), mice and rats. Endogenous p97 expression was observed in the hippocampus and substantia nigra pars compacta (SNpc) in mice and rats, the major pathological sites in Alzheimer’s^61^ and Parkinson’s disease respectively^62^ (Figure 3). Unfortunately, no tissue of NHP was available for other brain regions. Thus, p97 is endogenously expressed in regions of clinical interest that are affected by neurodegenerative diseases. Ultimately, the *segregase* and *unfoldase* activity of p97 in the brain could be advantageous for p97-PROTAC-based targeted degradation of toxic protein aggregates, especially for neurodegenerative diseases.

**Figure 3.**
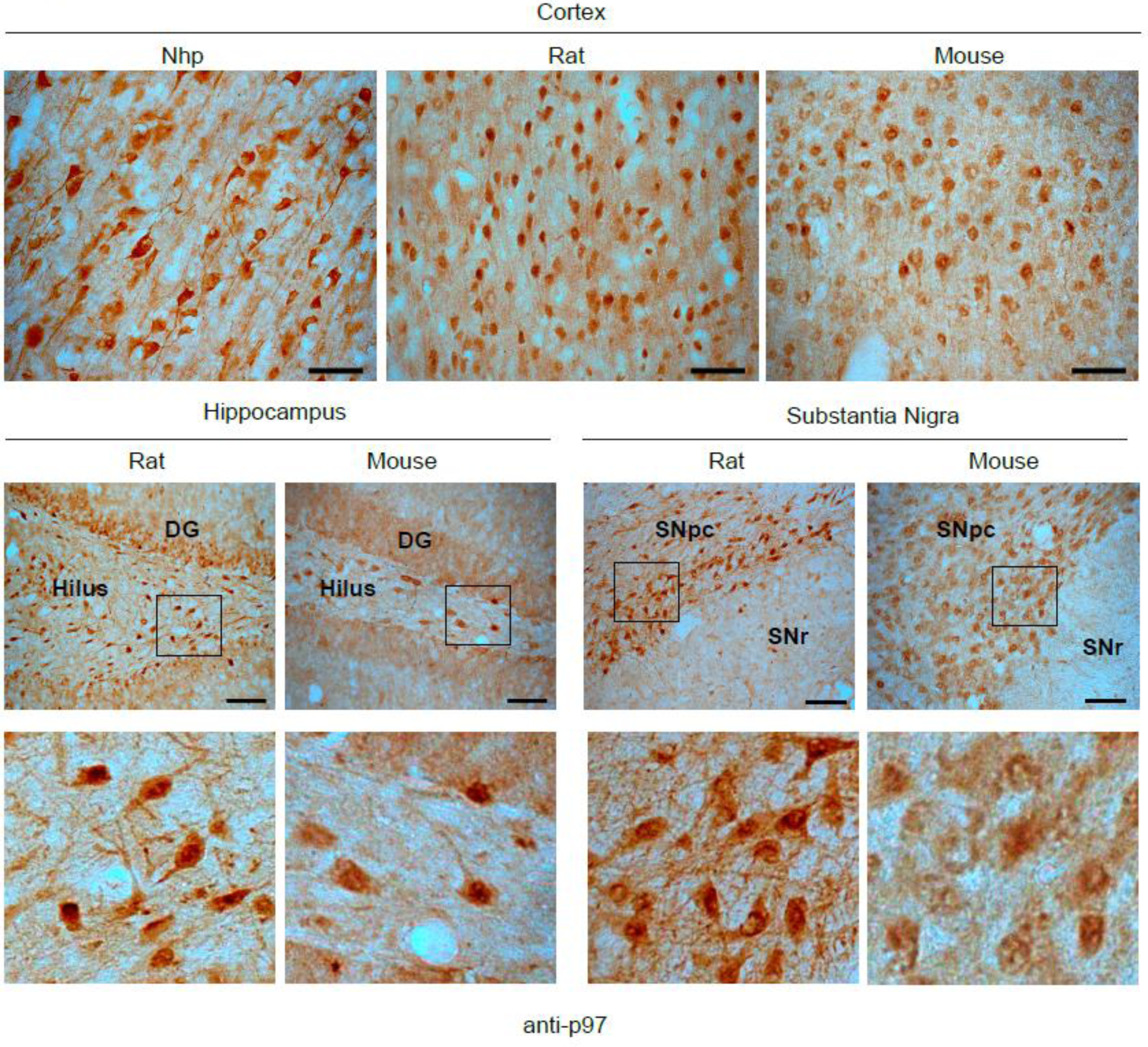
Endogenous p97expression in brain tissue. Immunohistochemistry against p97 in brain tissue sections from Non-Human Primates (Nhp) *Macaca fascicularis*, rat (*Sprague Dawley*) and mouse (C57BL6/C). p97 expression was detected in substantia nigra (SNpc), hippocampal, and cortical neurons. n=4

### Structural superposition of p97, Ubx, Nb^(GFP)^ and GFP

To gain insight into the structure of substrate-bound p97-PROTAC in the context of the p97 assembly, an AlphaFold2-based model of the p97-PROTAC/GFP-complex bound to p97 was generated (Figure 4A) (see Methods for details of the modelling procedure). After manually adjusting the conformation of the flexible linker between PROTAC Ubx and Nb^(GFP)^ moieties, the model shows that PROTAC-recruited GFP can access the central p97 pore (<15 Å distance between the pore entrance and the closest GFP residue) (Figure 4A & 4B). Consistent with the model, we determined that GFP alone, a barrel conformation protein of 28 kDa, can be recruited by the p97-PROTAC and is efficiently processed for degradation (Figure 1B, 4C & 4D). To test the mechanism by which the GFP became degraded, we co-transfected cells with GFP and either, an empty vector or with the p97-PROTAC Ubx-Nb^(GFP)^. After 20h, cells were treated with the proteasome inhibitor MG132 or DMSO as control. Upon MG132 treatment, we observed a full rescue of GFP protein levels. Therefore, by using a small substrate such as GFP as a model, we demonstrated that p97-PROTAC proteolytic activity relies on proteasome-mediated degradation (Figure 4E & 4F). Furthermore, upon depletion of p97 by siRNA transfection (Figure 4G), a significant rescue of GFP-Emerin protein levels was observed (Figure 4H), concomitant with an increase of the levels of the p97-PROTAC Ubx-Nb^(GFP)^ (Figure 4I), indicating that the PROTAC Ubx-Nb^(GFP)^ itself is also a substrate of the p97. To test the hypothesis that p97-PROTAC action is a ubiquitination-independent process, we globally inhibited ubiquitination using PYR41, a cell-permeable, irreversible inhibitor of the ubiquitin-activating enzyme E1 activity. First, we demonstrated that PYR41 was able to stabilize p53 in HeLa cells (Supplemental Figure 1D). Next, we used GFP-Emerin as a substrate model and treated the cell with either DMSO or 50uM of PYR41 for 4h, and in accordance with our hypothesis, no inhibition of GFP-Emerin degradation was observed by the PYR41 treatment (Figure 4J & 4K). We also investigated whether the activities of p97 were required for the substrate mediated degradation triggered by p97-PROTAC Ubx-Nb^(GFP)^. We tested the degradation of GFP-Emerin in HeLa cells treated with either DMSO or CB-5083, a potent inhibitor of the ATPase activity of p97. As an internal control, we measured the levels of CHOP, a sensor of ER stress known to be induced by p97 inhibitors (Figure 4L). Surprisingly we did not observe inhibition of degradation of GFP-Emerin in cells treated with CB-5083, despite a slight increase in CHOP levels (Figure 4M). Thus, our results show that p97-PROTAC targets proteins for proteasomal mediated degradation by a ubiquitin-independent mechanism.

**Figure 4.**
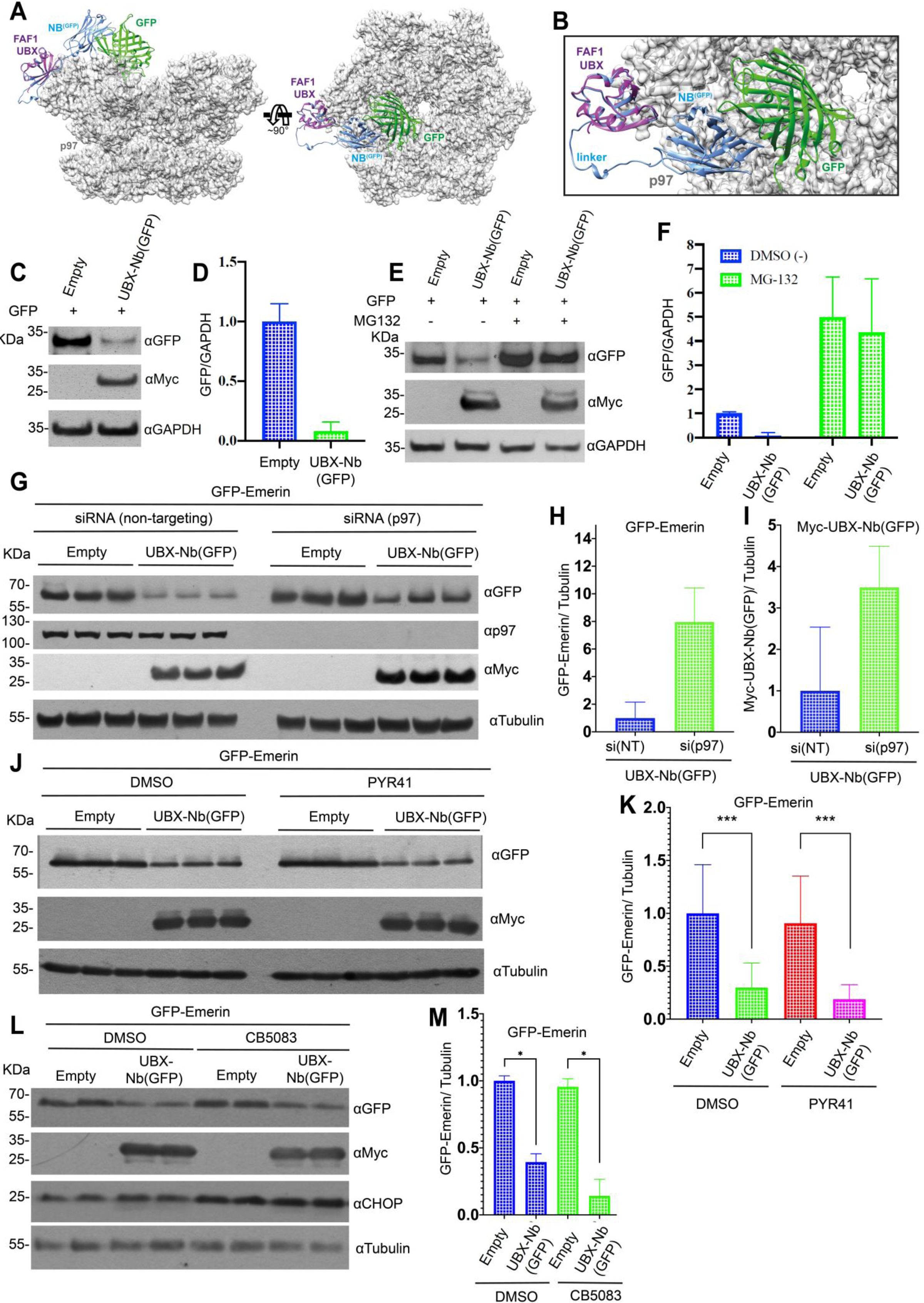
Molecular model and degradation activity of p97-PROTAC. **A)** Model representations of the FAF1 UBX domain (purple), UBX-Nb^(GFP)^ (blue), GFP (green), and the p97 hexamer (light grey cartoon with semi-transparent molecular surface representation). **B)** A magnified view of the model shown in A. **C)** GFP monomer was co-transfected with UBX-Nb ^(GFP)^ or empty vector in HeLa cells. Protein degradation was analyzed by western blot analysis. **D)** Quantification of C. **E)** GFP monomer was co-transfected with UBX-Nb^(GFP)^ or empty vector in HeLa cells, after 24 h were incubated with DMSO or the proteasome inhibitor MG132 (25 uM final concentration) for 4h. Protein degradation was analyzed by western blot. **F)** Quantification of E. **G)** p97 was silenced by transfection with p97-siRNA in HeLa cells. Subsequently, the cells were transfected with GFP-Emerin and either the empty vector or UBX-Nb^(GFP)^ vector. Protein degradation was analyzed western blot analysis. **H)** Quantification of GFP-Emerin in cells treated with the UBX-Nb^(GFP)^ vector in HeLa cells treated with either and siNT control or an SiRNA p97 siRNA. **I)** Quantification of the UBX-Nb^(GFP)^ in cells treated with the UBX-Nb^(GFP)^ vector in HeLa cells treated with either and siNT control or an SiRNA p97 **J)** HeLa cells co-transfected with GFP-Emerin and either empty vector or UBX-Nb^(GFP)^ in addition the cells were treated with the E1 ubiquitin inhibitor PYR-41 (50 µM) for 4 hours at 37°C. Subsequently, total proteins were extracted, and protein degradation was analyzed by western blot. **K)** Quantification of J. **L)** GFP-Emerin was co-transfected with UBX-Nb^(GFP)^ or empty vector in HeLa cells, after 24 h the cells were incubated with DMSO (as control) or the p97 inhibitor CB-5083 (4 uM final concentration) for 6h. Protein degradation was analyzed by western blot using total proteins. M) Quantification of L. Western blots were quantified and statistically analyzed using a student’s t-test. P < 0.05 compared to controls. n=3.

### Degradation of proteins of clinical interest by a p97-based PROTAC

A hallmark for several neurodegenerative diseases is the accumulation of toxic aggregates in the intra- or extracellular space^63^. PROTAC technology has the potential to become a new therapeutic approach for proteotoxic diseases^64^. High levels of ubiquitin are found in intracellular aggregates, suggesting that ubiquitination-mediated proteasome degradation may be partially impaired ^65,66^. Targeting p97 to toxic aggregates can potentially contribute to the clearance of aggregates by inducing segregation, unfolding, and proteasomal degradation simultaneously. We tested two aggregate models of clinical interest for p97-PROTAC-mediated degradation, Huntingtin and α-Synuclein. In the first experiment, we used a GFP-fusion plasmid containing the exon 1 of HTT with either 23 CAG repeats (GFP-HTT Q23, wild-type HTT) or 74 CAG repeats (GFP-HTT Q74, mutant HTT)^67^. We observed efficient degradation of GFP-HTT Q23 by the p97-PROTAC Ubx-Nb^(GFP)^ (Figure 5A & 5B) and a co-localization of p97-PROTAC Ubx-Nb^(GFP)^ with HTT GFP-HTT Q23 (Figure 5C). Subsequently, we repeated the experiment to test the degradation of GFP-HTT Q74 and observed aggregate formation after mutant HTT transfection concomitant with efficient degradation of GFP-HTT Q74 by the p97-PROTAC Ubx-Nb^(GFP)^ (Figure 5D-F).

**Figure 5.**
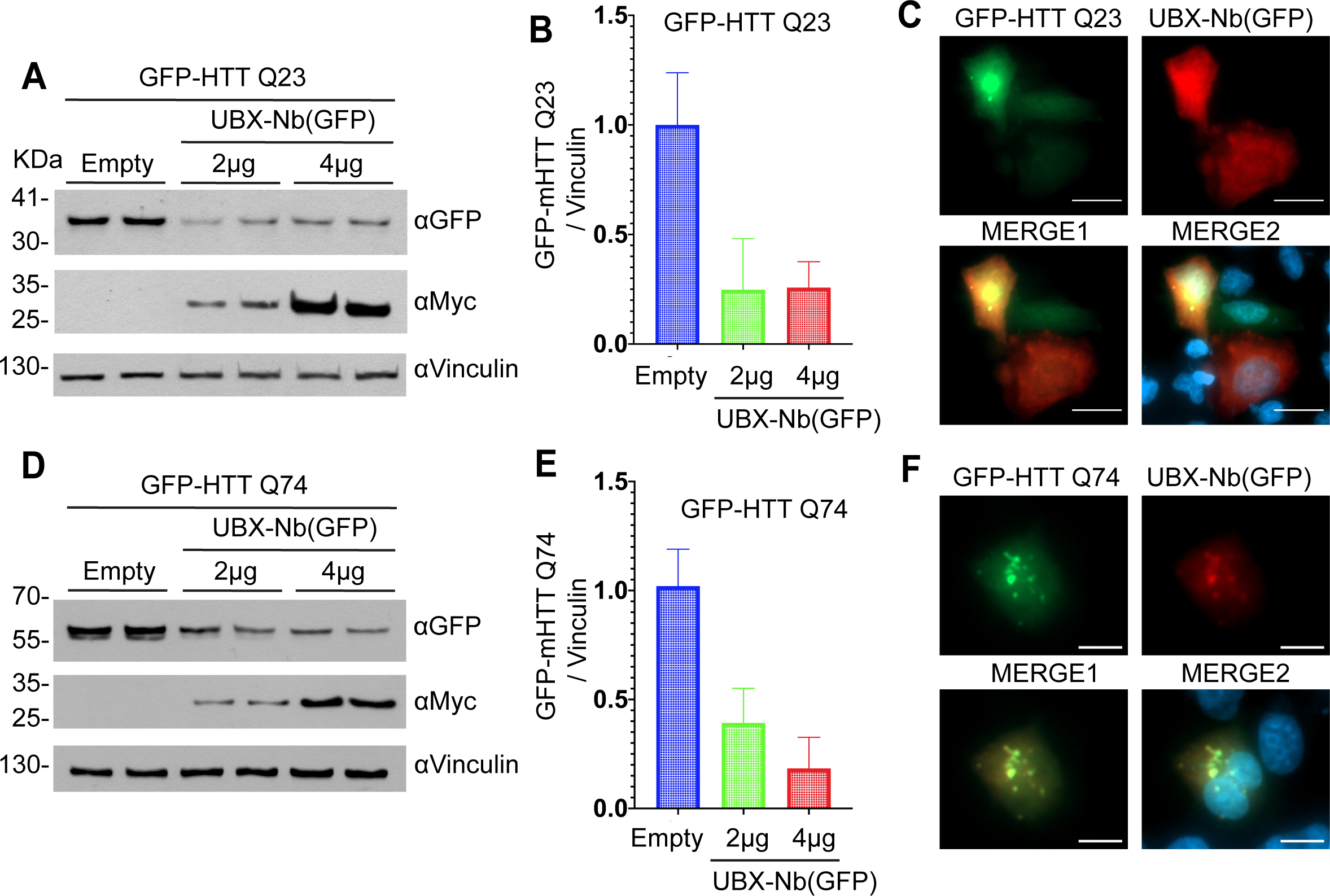
Degradation of Huntingtin wild type and mutant with p97-PROTAC UBX-Nb^(GFP)^. HeLa cells were transiently co-transfected with HTT GFP-tagged plasmids containing either 23 CAG repeats (GFP-HTT ^Q23^ - wild-type HTT) or 74 CAG repeats (GFP-HTT ^Q74^: mutant HTT) **A)** HeLa cells co-transfected with GFP-HTT ^Q23^ and increasing amount of the p97 PROTAC UBX-Nb^(GFP)^, degradation was determined by western blot analysis **B)** quantification of A; **C)** colocalization of GFP-HTT ^Q23^ and p97 PROTAC UBX-Nb^(GFP)^ **D)** HeLa cells were co-transfected with GFP-HTT ^Q74^ and increasing amount of the p97 PROTAC UBX-Nb^(GFP)^, degradation was determined by western blot analysis **E)** quantification of D; **F)** colocalization of GFP-HTT ^Q23^ and p97 PROTAC UBX-Nb^(GFP)^. Western blots were quantified and statistically analyzed using a student’s t-test. P < 0.05 compared to controls. n=3.

In the second challenge, we tested the capability of the p97-PROTAC Ubx-Nb^(GFP)^ to mediate the degradation of a GFP-tagged α-Synuclein mutant A53T and observed efficient degradation (Figure 6A & 6B). Then we generated another p97-PROTAC by replacing the anti-GFP nanobody with a specific α-Synuclein nanobody (NbSyn87)^68-71^. Likewise, cells were transfected with an untagged α-Synuclein A53T mutant together with an empty vector (as control) or with increasing concentrations of the anti-α-Synuclein p97-PROTAC Ubx-Nb^(Syn87)^. Again, we observed efficient degradation of the α-Synuclein A53T p97-PROTAC (Figure 6C & 6D). Thus, we demonstrated that the nanobody component on the p97-PROTAC is exchangeable and that the p97-PROTAC system is suitable for degrading human proteins of clinical interest.

**Figure 6.**
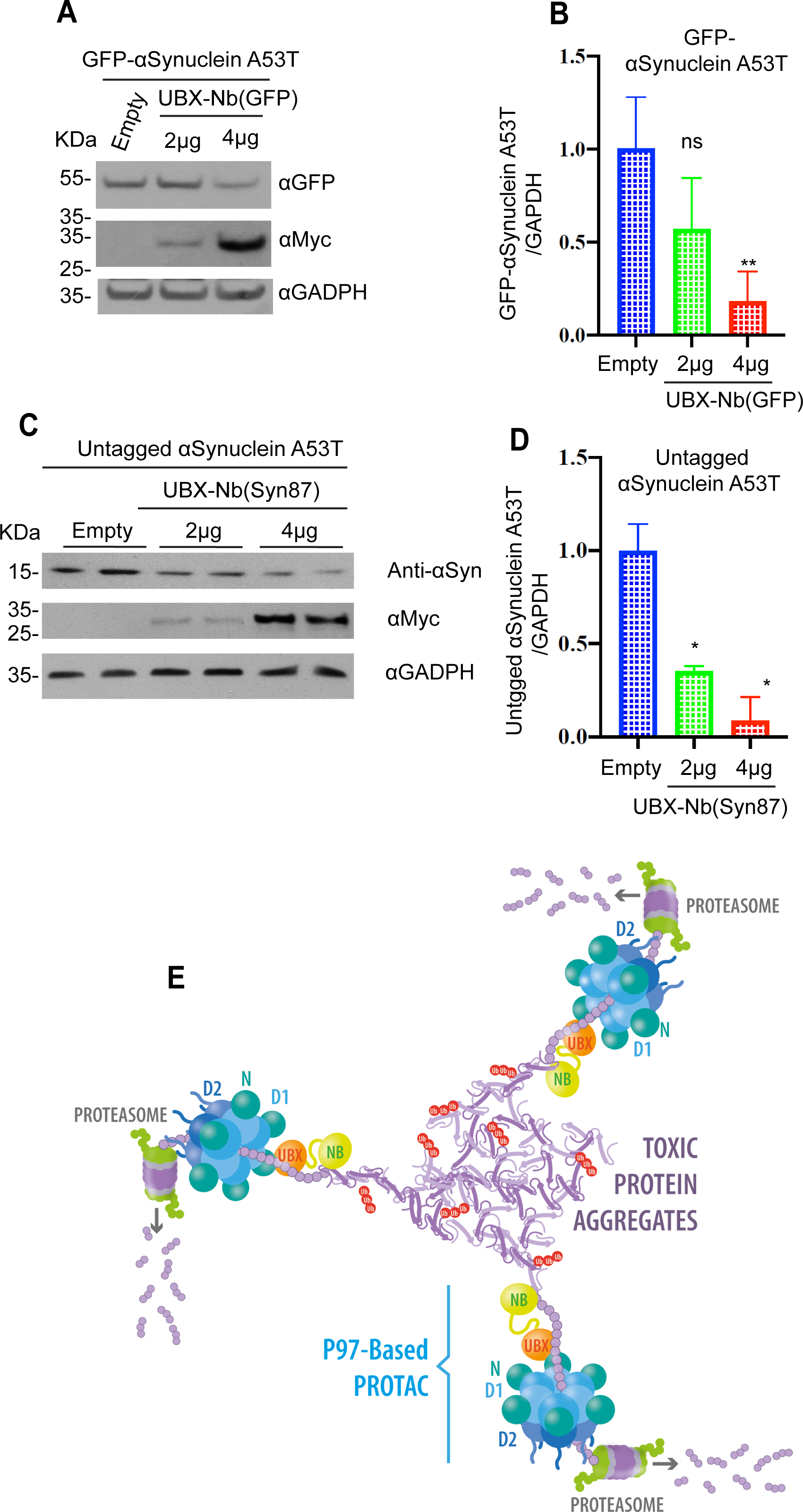
Degradation of Alpha Synuclein with p97-PROTAC. **A)** HeLa cells were co-transfected with a vector expressing αSynuclein mutant A53T fused to GFP (GFP-αSynuclein A53T) and empty or increasing concentrations of UBX-Nb^(GFP)^. A53T-GFP degradation was determined by Western blot. **B)** Quantification of A. **C)** Cells were co-transfected with a vector expressing untagged alpha synuclein mutant A53T and an empty vector or increasing concentrations of UBX-Nb^(Syn87)^. Untagged αSynuclein A53T degradation was determined by Western blot using an anti αSynuclein antibody**. D)** Quantification of C. **E)** Representative model of p97-PROTAC functioning in the degradation of proteins and protein aggregates. Western blots were quantified and statistically analyzed using a student’s t-test. P < 0.05 compared to controls. n=3.

## Discussion

The constitutive degradation of proteins is a rapid negative-regulatory mechanism, often involved in important stress response pathways. For example, under normoxia, Hif1α is rapidly hydroxylated in an oxygen-dependent manner, and the hydroxylation is recognized by an E3 Ligase receptor called VHL leading to continuous ubiquitin-mediated proteasomal degradation^73,75-77^. Hypoxia decreases Hif1α hydroxylation, and consequently, Hif1α dissociates from VHL, resulting in rapid stabilization, accumulation, and activation of the hypoxia response. Therefore, endogenous degradation is a natural and efficient way to downregulate the amount and function of proteins.

PROTAC technologies today rely on ubiquitin-mediated proteasomal degradation, but abnormal protein aggregation has multiple detrimental effects in the ubiquitin-mediated proteasomal degradation, including a significant reduction of the free ubiquitin pool^78^ and abundant ubiquitin accumulation into proteotoxic aggregates. Our p97-PROTAC technology provides an alternative to ubiquitin-mediated degradation.

We showed that an engineered p97-PROTAC consisting of a Ubx domain fused to a camelid nanobody was sufficient to trigger specific protein degradation in cells and further, using the ubiquitin E1 inhibitor PYR41, we demonstrated that p97-PROTACs are not inhibited by ablation of ubiquitination. In addition, using siRNA specific against p97, we inhibited the degradation of the substrate model. Intriguingly, we could not inhibit the action of p97-PROTACs with the specific p97 ATPase inhibitor CB-5083. Thus, mechanistically we cannot distinguish if the p97-PROTACs overcame the effect of the inhibitor or if the degradation mediated by the p97-PROTACs is mediated only by induced proximity to the proteasome and independent of p97’s ATPase activity. Further experiments need to be performed to understand the mechanism by which p97-PROTACs mediate ubiquitin-independent degradation in cells.

p97 is an extensively characterized ubiquitously expressed protein involved in fundamental cellular processes, such as the degradation of proteins associated with the endoplasmic reticulum (ER) (ERAD)^79^, autophagy^80^, and it is also an important player in the proteostasis of aggregates in Parkinson’s disease^46,81,82^. Indeed, p97 has been shown to protect against the proteopathic spread of pathogenic aggregates in animal models^83^. In this study, we characterized p97 expression in the brains of animal models and observed ubiquitous p97 expression in all sections. Specifically, high p97 expression levels were detected in the cortex, substantia nigra pars compacta and hippocampus, regions highly prone to protein aggregation and neurodegeneration in neurodegenerative diseases such as Parkinson’s and Alzheimer’s disease^63^. These observations support the possibility of using p97-based PROTACs to reduce protein aggregation in neurodegeneration-related targets in the future.

The PROTAC technologies might increase the number of druggable proteins by a change in the inhibition paradigm. In contrast to conventional drugs that aim for chemical modulation of the enzymatic activity or function, PROTAC aims at the downregulation of the levels of specific targets based exclusively on selective binding and degradation.

In conclusion, our work unveils the potential of a p97-PROTAC as the first E3 ubiquitin ligase-independent technology that targets proteins for degradation at diverse subcellular locations, integral membrane protein residing at the inner nuclear membrane, chromatin located and liquid– liquid phase separated compartments, also providing a new technology to target protein aggregates of clinical interest for proteasomal degradation.

## Material and Methods

### Design and cloning

The constructs consist mainly of three domains: an anti-GFP nanobody^20^, which binds to the target protein; a linker and the UBX domain of the p97 Adapter FAF1. The linker used to connect both domains, KESGSVSSEQLAQFRSLD, was originally designed for the construction of single-chain antigen-binding proteins, connecting its constituent domains with sufficient flexibility to avoid hindering its antigen-binding capacity^85^. Anti-GFP nanobody can recognize both, the YFP and GFP protein.^57^ In addition, a myc-tag was added to prove expression. The full sequence was cloned into the pcDNA 5 FRT/TO vector by the Gibson Assembly technique. Later, the anti-GFP nanobody was cut with restriction enzymes to be replaced by an anti-α-Synuclein (Nb87) nanobody NbSyn87^68-71^.

### Cell culture

All cells (HeLa and U2OS) were cultured in Dulbecco’s Modified Eagle’s medium (DMEM; Gibco) supplemented with 10% fetal bovine serum and 100 units/mL of penicillin and streptomycin and maintained at 37°C in a humidified incubator at 5% CO2. Plasmid transfection was performed in 6-well plates using 4 μg of DNA, 24 h after transfection, cells were lysed, and proteins were collected. Cells were transiently transfected with the following vectors: pcDNA5 FRT/TO GFP-ETV1, pcDNA5 FRT/TO GFP-Emerin, pEYFP-Coilin, pcDNA FRT/TO GFP, pEGFP-C1-tagged plasmids containing the exon 1 of HTT with 23 CAG repeats (GFP-Q23: wild-type HTT. Addgene, #40261) or 74 CAG repeats (GFP-Q74: mutant HTT. Addgene, #40262). Vectors to alpha synuclein were a gift from our collaborator Dr. Gopal Sapkota (pcDNA5-FRT/TO-GFP-SCNA-A53T, #59047 or pcDNA5-FRT/TO-SCNA-A53T, #59042). As control, we used an empty vector pcDNA5 FRT/TO. Transfection was performed using Lipofectamine 2000 (Invitrogen) according to the manufacturer’s instructions, and media were supplemented with normocin (100 ug/mL) during transfection (Invivogen). For lysis, cells were washed twice in ice-cold phosphate buffer saline (PBS), scraped on ice in lysis buffer (Tris–HCl pH 6.8, NaCl, glycerol and SDS 10%), supplemented with complete protease inhibitors (one tablet per 25 mL: Roche), and 0.1% b-mercaptoethanol (Sigma). Cell extracts were either cleared and processed immediately or stored at -20°C. The protein concentration was determined in a 96-well format using the Pierce BCA protein assay kit (Thermo Fisher Scientific).

### Antibodies & Inhibitors

We used the following primary antibodies; anti-GFP (Invitrogen, GF28R mAb MA5-15256), anti-α-Synuclein (Santa Cruz Biotechnology, 3H2897 mAb sc-69977), anti-53BP1 (Invitrogen, 53BP1 Polyclonal Antibody PA1-16565), anti-Myc-tag (Cell Signaling, 9B11 Mouse mAb #2276), anti-vcp/p97 (Sigma-Aldrich Polyclonal Antibody HPA012728), anti-p53 (Invitrogen, monoclonal antibody DO-7 # MA5-12557), anti-CHOP (Cell Signaling, L63F7 Mouse mAb #2895), anti-ubiquitin (Sigma-Aldrich ST1200 Mouse mAb FK2), anti-alpha Tubulin (Santa Cruz Biotechnology, B7 mAb sc-5286), anti-GAPDH (Santa Cruz Biotechnology, mAb sc-47724), anti-vinculin (Santa Cruz Biotechnology, 7F9 mAb sc-73614). Horseradish peroxidase (HRP)-coupled secondary antibodies and Alexa Fluor secondary antibodies were purchased from Invitrogen (ThermoFisher): Mouse IgG, IgM (H+L) Secondary Antibody (A-10677), Rabbit IgG (H+L) Secondary Antibody (31460) and Mouse IgG (H+L) Highly Cross-Adsorbed Secondary Antibody (A-11032). The inhibitors used in this work were: MG132, a proteasome inhibitor from Sigma-Aldrich (catalog number 474790); PYR-41, an E1 ligase inhibitor from Sigma-Aldrich (catalog number N2915); and CB-5085, a D2 domain inhibitor of p97 from Cayman Chemical.

### Generation of a 53BP1 endogenously tagged YFP-53BP1 using CRISPR/Cas9

U2OS T-Rex cells were co-transfected with the pOG44 plasmid, which constitutively expresses the Flp recombinase and pcDNA5 FRT/TO codon optimized Streptococcus pyogenes M1 GAS Cas9 D10A-NLS-FLAG (DU45732 MRC-PPU reagent) for isogenic integration of the cassette. To decrease possible off-target effects, we applied the CAS9 D10A nickase system for genetic cleavage. Single clones were selected to generate the U2OS stable expressing Cas9 D10A cells, U2OS SEC-C D10A. Two specific guide RNA (gRNA) targeting sequences were identified in the 5’UTR before the 53BP1 start codon using the E-CRISPR software (http://www.e-crisp.org/E-CRISP/reannotate_crispr.html). The forward gRNA g53BP1-1 5’AGACCTCTAGCTCGAGCGCGAGG 3’ and a reverse gRNA g53BP1-7 5’GTCCCTCCAGATCGATCCCTAGG 3’ were cloned into pU6-Puro via site-directed mutagenesis using the QuickChange method (Stratagene) cloned using the previously described methodology^86,87^ and confirmed by DNA sequencing. We designed a strategy for YFP delivery at the N-terminus of the 53BP1 protein using a CRISPR/Cas9 knock-in methodology. In short, the sense and antisense gRNAs were transfected in U2OS T-Rex cells and modified to produce CAS9 D10A nickase in a doxycycline-inducible manner. We also engineered a synthetic vector with two homologous flanking regions around the cleavage site, where we inserted YFP cDNA and small linker in front of the endogenous 53BP1 gene synthetically produced by GeneArt (Life Technologies) (supplemental methods). The gRNA recognition sites were mutated on the synthetic donor to avoid cleavage by the gRNA/Cas9 complex. Finally, the two gRNAs and the donor vector carrying the YFP cDNA flanked with the homologue regions were transfected in U2OS cells. The day after the cells were transfected, the expression of the Cas9 nuclease was induced by adding 1 µg/mL doxycycline and the cells were incubated for 4 days before FACS analysis to identify YFP positive cells and further cell sorting based on single cell isolation.

### Flow cytometry and cell sorting

Cells were analyzed for YFP fluorescence on an LSR Fortessa or FACS Canto flow cytometer (Becton Dickinson) and data were analyzed using FlowJo software (Tree Star Inc). Single cells were identified based on FSC-A, FSC-W and SSC-A, and YFP fluorescence measured with 488 nm excitation and emission detected at 530±30 nm. Cell sorting was performed on an Influx cell sorter (Becton Dickinson) with FACS Software, using the same cell identification procedure described above. Single YFP expressing cells were sorted onto individual wells of a 96-well plate containing DMEM supplemented with 20% fetal bovine serum (FBS), 2 mM L-glutamine, 100 units/ml penicillin, 100 μg/ml streptomycin, and 100 µg/ml transfection compatible antibiotic normocin 1x (Invivogene). Single-cell clones were left to proliferate, and YFP fluorescence was determined.

### Immunofluorescence and high content microscopy

HeLa cells transfected with GFP proteins were grown in a 96-well optical plate (Thermo Fisher Scientific) or 24-well lidded plates. Cells were fixed with 4% paraformaldehyde at 37°C for 10 min. Cells were washed with 1x PBS and permeabilized in 0.2% TritonX100 PBS. After washing the cells three times in 1x PBS, they were incubated with blocking solution (FBS 5%-PBS 1x) for 30 minutes and then incubated with the primary antibody for 1 hour at 37 ° C. After washing another three times with 1x PBS, a secondary antibody Alexa Fluor was used at 1: 3000, it was incubated for 45 min at 37 ° C. For nuclei staining, cells were washed with PBS and incubated for 10 min at room temperature with 0.1 mg / ml DAPI. After the final wash, the cells were kept in the 96-well optical plates in 1x PBS. Cover-fixed cells were mounted on slides. Fixed cell images were acquired with a high content automated microscope, Celldiscoverer 7 (Carl Zeiss GmbH, Jena, Germany).

### Silencing assay

HeLa cells were cultured in a 6-well plate and transfected with siRNAs control siRNA-B, (sc-44230), or p97 siRNA (sc-37187) at 40% confluence. Transfection was performed using Lipofectamine RNAiMax (Invitrogen) according to the manufacturer’s instructions, and the media were supplemented with Normocin (Invivogen) during transfection. Twenty-four hours post siRNA transfection, the medium was changed to fresh DMEM-Normocin, and GFP-Emerin, as well as both the empty vector and the Ubx-Nb (GFP) vector, were co-transfected. The complexes were left in culture for 48 hours. Subsequently, total proteins were extracted with lysis buffer to perform the western blot assay.

### SDS–PAGE and western blotting

Reduced protein extracts were separated on 10% SDS–PAGE gels or 4–12% NuPAGE bis–tris precast gels (Invitrogen) by electrophoresis. Proteins were then transferred onto nitrocellulose membranes (Millipore). Non-specific sites were blocked with blocking solution (PBS containing 0.1% Tween20 with 5% (w/v) non-fat milk) for 30 min at room temperature with agitation. The blocking solution was discarded, and membranes were incubated overnight at 4°C in 5% BSA PBS-T (PBS 1x containing 0.1% Tween20) with the appropriate primary antibodies. Membranes were then washed in PBS-T and incubated with HRP-conjugated secondary antibodies in 5% milk-PBS-T for 1 hour at room temperature, followed by 3 x 5 min washes with PBS-T and visualized using the ECL reagent (Pierce) and exposure on medical X-ray films.

### Animals, fixation, and tissue processing

Mouse (C57BL6 / C mice), rat (sprague dawley), and monkey (Macaca fascicularis) brain sections were obtained from the brain bank of HM CINAC. All procedures with animals were carried out in accordance with the Directive of the Council of the European Communities (2010/63 / EU) and Spanish legislation (RD53 / 2013) on animal experimentation^88^. Animal perfusion, fixation, and tissue processing was carried out as previously described^88,89^.

### Brain immunohistochemistry

Coronal free-floating 30-mm thick sections were washed with Tris buffer (TB) and treated with citrate buffer (pH 6) for 30 min at 37° C for antigen retrival. Inhibition of endogenous peroxidase activity was carried out using a mixture of 10% methanol and 3% concentrated H_2_O_2_ for 20 min at room temperature. Normal serum of the same species as the secondary antibody (Normal Goat Serum, NGS) was applied for 3 h to block non-specific binding sites. The sections were immunostained with anti-p97 antibody (Rabbit polyclonal anti-vcp, sigma aldrich HPA012728, dilution 1: 1000) at 4 ° C for 72h. The sections were washed with Tris-buffered saline (TBS) and incubated for 2 h with a solution containing the secondary biotinylated antibody (goat anti-rabbit, dilution 1:400; Chemicon, Burlington, USA) in TBS-NGS. The sections were then incubated for 45 min with the avidin-biotin-peroxidase complex (PK-6100, ABC Vectastain; Vector Laboratories, Burlingame, USA). Immunohistochemical reactions were visualized by incubating the sections with 0.05% 3,3-diaminobenzidine (DAB, Sigma, St. Louis, USA) and 0.003% H_2_O_2_. The sections were then dehydrated with graduated ethyl alcohol (EtOH) and rinsed in two xylene changes before being mounted in dibutyl polystyrene xylene phthalate (DPX) and applying glass coverslips. Omission of the primary antibody resulted in a lack of staining (images not shown).

### Molecular modelling

An atomic model of Ubx-Nb^(GFP)^ was predicted using the AlphaFold2 algorithm via the ColabFold notebook^90,91^.The FAF1 UBX domain position on the p97 hexamer was modelled by LSQ superposition of the p97 N-domain portion of a p97-N/FAF1 UBX co-crystal structure (PDB 3qq8)^92^ with the N-domain of chain A of a p97 hexamer cryo-EM structure (PDB 7jy5)^93^ using the program Coot^94^. Subsequently, the FAF1 UBX domain portion of the AlphaFold prediction was aligned with the p97/FAF1 UBX domain model using LSQ superpose in Coot. The conformation of the flexible linker between Nb^(GFP)^ and FAF1 UBX domain in the fusion protein model was manually adjusted using the “Regularize Zone” feature in Coot. Finally, the position of GFP was modelled by LSQ superposition of the nanobody portion of the adjusted Ubx-Nb^(GFP)^ coordinates with the crystal structure of a Nb^(GFP)^/GFP co-crystal structure (PDB 3ogo)^57^.

### Statistical Analysis

The results are expressed as the mean ± SEM. All statistical analyses were performed using the GraphPad Prism software. For all western blots, an unpaired student’s t-test was performed to determine differences between control cells with an empty vector versus Ubx treated cells. A confidence level of 95% was accepted as significant.

### Author Contributions Statement

Conceptualization: CSR, PCC and ARF; cloning and design: CSR, GVN, GPS and ARF; animal tissue acquisition, staining, and analysis: CSR, JB, NLGR, MMR, LFB, PCC and JO; immunofluorescence and microscopy images: CSR, JB and ARF; CRISPR/Cas9 editing ARF; biochemical studies and western blotting: C.S.R, structural modelling: D.S; analysis: CSR, LFB and ARF; funding acquisition PCC, JO and ARF; resources: PCC, JO and ARF; supervision: ARF; writing – original draft: CSR, PCC and ARF. All authors have contributed to writing, reviewing, and editing and have agreed to the published version of the manuscript.

## Acknowledgments

This work was funded by FONDECYT REGULAR 1200427 to ARF; FONIS EU-LAC T010047 to CSR, PCC, JO, JB, NLR & ARF; to ARF; ANID-EXPLORACIÓN 13220075 to ARF & GVN; ANID 3220635 to GVN; MPG 190011 to ARF; ANID-STINT CS2018-7952. J.B was funded by ISCIII Miguel Servet Program CP19/00200 and FIS PI20/00496. Graduate fellowship ANID N° 22170632 to C.S; N.L.G.R was funded by Becas Santander Iberoamerica Investigacion 2018/2019 and grant S2017/BMD-3700 (NEUROMETAB-CM) from Comunidad de Madrid; D.S. was supported by the German Research Foundation (DFG) Emmy Noether Programme SCHW1851/1-1. We would like to thank Anne Berking and Amber Philp for proofreading and Felipe Serrano for illustration.

