## Supplemental Figures for "Specific proteolysis mediated by a p97-directed proteolysis-targeting chimera (PROTAC)"

### Supplemental Figure-1

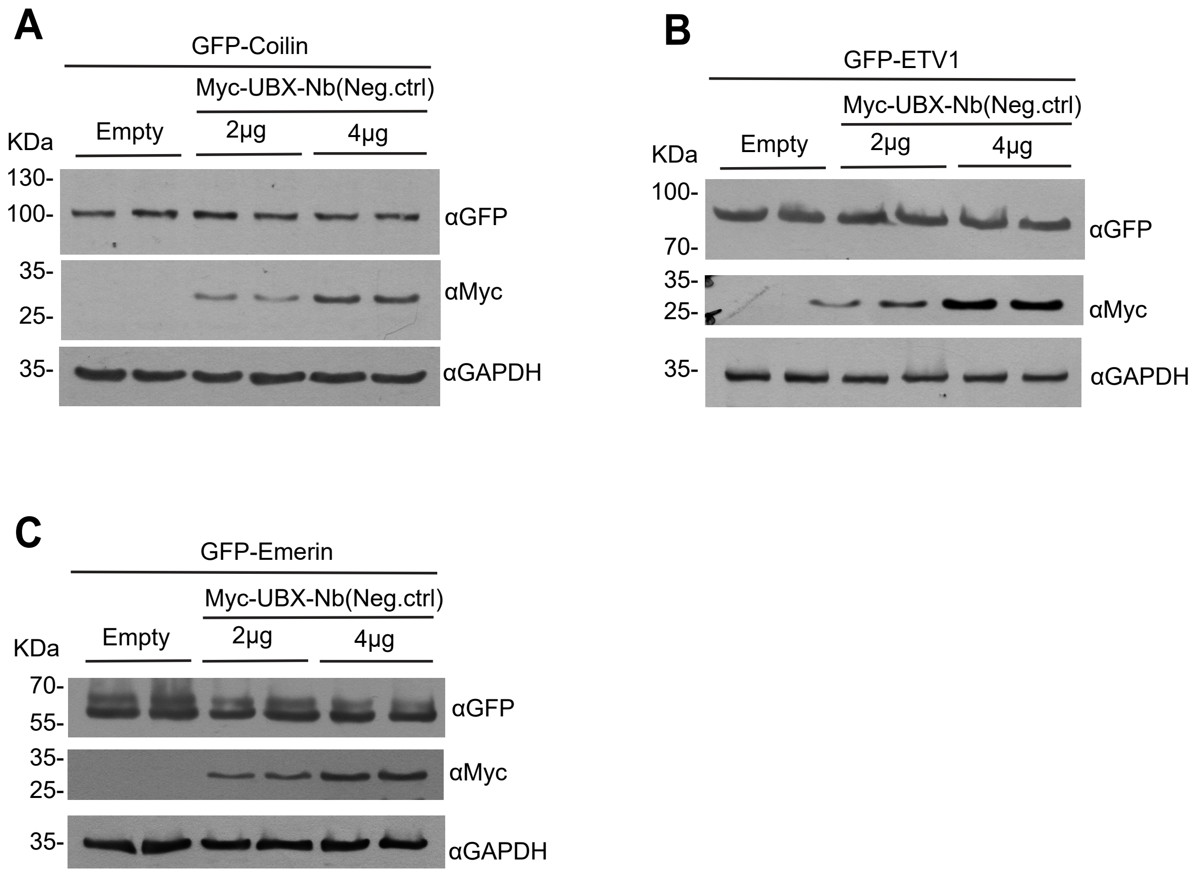


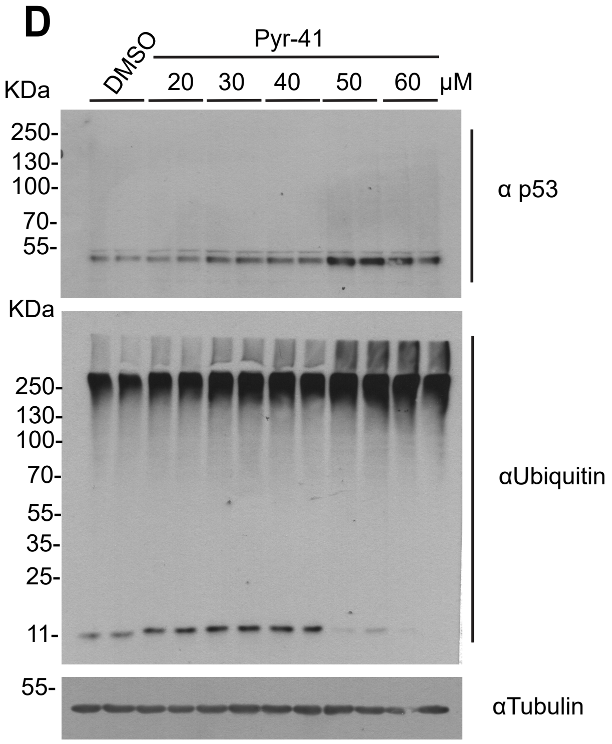


Figure 1–figure supplement 1.- A – C) HeLa cells were co-transfected with GFP-Coilin (A), GFP-ETV1 (B) or GFP-Emerin (C) with and empty vector or increasing concentrations of UBX-Nb(Neg.ctrl) vector. Protein degradation was analyzed by Western blot. D) HeLa cells were incubated with different concentrations of PYR-41 (a cell permeable irreversible inhibitor of ubiquitin-activating enzyme E1) for 4 hours. Subsequently, total cellular proteins were extracted, and Western blot analyses were performed. Nitrocellulose membranes were incubated with the primary antibodies anti-p53 and anti-ubiquitin antibody. DMSO was used as a control. n=2
